# Motor unit recruitment patterns of the quadriceps differ between continuous high- and low-torque isometric knee extension to momentary failure

**DOI:** 10.1101/2021.04.08.438966

**Authors:** Jonathan Murphy, Emma Hodson-Tole, Andrew D. Vigotsky, Jim R. Potvin, James P. Fisher, James Steele

## Abstract

The size principle is a theory of motor unit (MU) recruitment that suggests MUs are recruited in an orderly manner from the smallest (lower threshold) to the largest (higher threshold) MUs. A consequence of this biophysical theory is that, for isometric contractions, recruitment is dependent on the intensity of actual effort required to meet task demands. This concept has been supported by modelling work demonstrating that, in tasks performed to momentary failure, full MU recruitment will have occurred upon reaching failure irrespective of the force requirements of the task. However, *in vivo* studies examining this are limited. Therefore, the aim of the current study was to examine MU recruitment of the quadriceps under both higher- and lower-torque (70% and 30% of MVC, respectively) isometric knee extension, performed to momentary failure. Specifically, we compared surface electromyography (sEMG) frequency characteristics, determined by wavelet analysis, across the two continuous isometric knee extension tasks to identify potential differences in recruitment patterns. A convenience sample of 10 recreationally active adult males (height: 179.6±6.0 cm; mass: 76.8±7.3 kg; age: 26±7 years) with previous resistance training experience (6±3 years) were recruited. Using a within-session, repeated-measures, randomised crossover design participants performed the knee extension tasks whilst sEMG was collected from the vastus medialis (VM), rectus femoris (RF) and vastus lateralis (VL). Myoelectric signals were decomposed into intensities as a function of time and frequency using an EMG-specific wavelet transformation. Our first analysis compared the mean frequency at momentary failure; second, we investigated the effects of load on relative changes in wavelet intensities; finally, we quantified the degree of wavelet similarity over time. Wavelet-based calculation of the mean signal frequency appeared to show similar mean frequency characteristics occurring when reaching momentary failure. However, individual wavelets revealed that different changes in frequency components occurred between the two tasks, suggesting that patterns of recruitment differed. Low-torque conditions resulted in an increase in intensity of all frequency components across the trials for each muscle whereas high-torque conditions resulted in a wider range of frequency components contained within the myoelectric signals at the beginning of the trials. However, as the low-torque trial neared momentary failure there was an increased agreement between conditions across wavelets. Our results corroborate modelling studies as well as recent biopsy evidence, suggesting overall MU recruitment may largely be similar for isometric tasks performed to momentary failure with the highest threshold MUs likely recruited, despite being achieved with differences in the pattern of recruitment over time utilised.

## Introduction

The size principle is a theory of MU recruitment that suggests MUs are recruited in an orderly manner from the smallest (lower threshold) to the largest (higher threshold) MUs (Denny-Brown and Pennybacker, 1938; Henneman, 1957) as excitation is increased. A consequence of this biophysical theory is that recruitment is dependent on the intensity of actual effort required to meet task demands (Carpinelli, 2008). Therefore, in an exercise task performed to momentary failure, it is argued that an equivocal population of MUs may be recruited during protocols with both high- and low-loads (Carpinelli, 2008; Fisher et al., 2017). This concept has been supported by modelling work demonstrating that, in tasks performed to momentary failure, full MU recruitment will have occurred upon reaching failure irrespective of the force requirements of the task (Potvin and Fuglevand, 2017). However, *in vivo* studies examining this are limited.

The characteristics of human MUs have been examined for some time using electromyography (EMG) to measure the electrical activity in the muscles (Hodson-Tole and Wakeling, 2009; Duchateau and Enoka, 2011). Likely due to its ease of use, surface EMG (sEMG) was used in several recent studies (Schoenfeld et al., 2014; Jenkins et al., 2015; Looney et al., 2015; Schoenfeld et al., 2016; Gonzalez et al., 2017; Chapman et al., 2019) in which the aim was to investigate MU recruitment during exercise tasks under different loading schemes, in attempts to test the hypothesis of similar MU recruitment. Specifically, these prior studies compared sEMG amplitudes between higher- and lower-load resistance exercise conditions and found that both mean and peak sEMG amplitudes are higher when performed using higher loads (Schoenfeld et al., 2014b; Jenkins et al., 2015; Looney et al., 2015; Schoenfeld et al., 2016; Gonzalez et al., 2017). These findings led authors to the interpretation that greater MU recruitment occurs under high load conditions. Although, in some instances, peak amplitudes—particularly at the point of momentary failure—appear similar (Schoenfeld et al., 2016; Gonzalez et al., 2017; Chapman et al., 2019). Notwithstanding these findings, some argue that inferences regarding MU recruitment from simple amplitude-based analyses of sEMG are specious (Enoka & Duchateau, 2015; Fisher et al., 2017; Vigotsky et al., 2018).

Recent work shows that sEMG amplitude, and even median frequency characteristics, are poorly associated to MU recruitment (examined using high density multi-channel electrode arrays) at a range of force requirements (Del Vecchio et al., 2017). As such, others used alternative measurements to assess MU recruitment characteristics. Biopsy work lends contradictory evidence to the conclusions drawn from previous studies: low- and high-load tasks performed to momentary failure result in similar glycogen depletion despite disparate sEMG amplitudes (Morton et al., 2019). Thus, the differential sEMG amplitudes observed across loading conditions performed to momentary failure may, in fact, simply reflect different MU recruitment patterns or signal distortions, while total MU pool recruitment may still be similar across the tasks (Fisher et al., 2017; Vigotsky et al., 2017; Vigotsky et al. 2018).

There are several potential explanations for why differential sEMG amplitudes can arise with similar total MU recruitment. First, under higher load (and thus higher force/torque) conditions, a greater number of MUs need to be recruited synchronously (both higher and lower threshold MUs) and at increased firing rates in order to produce sufficient force. This highly ‘synchronous’ MU recruitment might be expected to result in greater sEMG amplitudes. Conversely, a sustained task using a lower load (and thus lower force/torque) might be expected to initially only recruit sufficient MUs to produce the necessary force (predominantly lower threshold MUs); however, as the task continues, previously recruited MUs fatigue and other MUs would need to be recruited to sustain the required force. Thus, at a given instance in time, relative to momentary failure during a sufficiently lower load condition, it might be expected that the number of MUs being recruited would be fewer than during higher load conditions, resulting in a lower sEMG amplitude. Second, sEMG amplitudes may also be constrained by changes in MU action potential shape at momentary failure under submaximal conditions, meaning that the observed amplitudes reflect changes in the peripheral environment rather than just neural drive (Dimitrova and Dimitrov, 2003; Dideriksen et al., 2011).

Third, ‘sequential’ MU recruitment, from lower threshold to higher threshold MUs, during fatiguing contractions may be permitted by reductions in recruitment thresholds (Adam and DeLuca, 2003; Contessa et al., 2016), suggesting that as momentary failure is neared the number of MUs recruited may be similar between higher and lower load conditions. Fourth, it has also been argued MUs may also ‘cycle’ to maintain force requirements (Jensen et al., 2000; Westad et al., 2003; Bawa et al., 2006). Under such circumstances, the total number of MUs recruited at any given point might be lower during lower load conditions, but instantaneous recruitment may not be indicative of recruitment across the duration of a trial. Fifth, firing rate adapts to the greatest degree in higher threshold MUs, which may further reduce measured EMG amplitudes even at momentary failure with all MUs recruited (Sawczuk et al., 1995; Gorman et al., 2005; Revill & Fuglevand, 2011), particularly under lower load conditions where the longer durations allow for more pronounced adapted decreases in firing rates (Potvin & Fuglevand, 2017). If performed to momentary failure it might be expected that eventually all available MUs would have been recruited in either higher- or lower-load conditions (Potvin & Fuglevand, 2017); however, potentially differing patterns of recruitment may occur.

The exact MU recruitment patterns during high- and low-load fatiguing tasks remain somewhat elusive (Fisher et al., 2017). Mammalian skeletal muscle is composed of a range of different fibre types (Schiaffino and Reggiani, 1994) that generate a range of myoelectrical properties during muscle activation (Wakeling and Syme, 2002). To examine MU recruitment during exercise to momentary failure, techniques more advanced than simple amplitude and frequency analyses may be required, such as spike-triggered averaging (Boe et al., 2004) or decomposition using high-density multi-channel electrode arrays (Del Vecchio et al., 2017). Indeed, a recent study by Muddle et al., (2018) used decomposition sEMG to examine MU recruitment and firing behaviours of the vastus lateralis during repeated isometric knee extension to momentary failure with both higher- and lower- torque conditions. They reported that, although both conditions resulted in recruitment of additional higher threshold MUs to maintain torque production, on average, there was greater recruitment of larger MUs in the higher-torque condition. To our knowledge, this is the first study to have examined MU behaviour under different torque requirements using such a technique. Though, another recent study (Harmon et al., 2021) has also examined motor unit action potentials (MUAPs) using decomposition sEMG in non-fatiguing high-torque and fatiguing low-torque conditions performed to momentary failure, suggesting similar MUAPs between conditions when the latter reached momentary failure. However, it is acknowledged that there is debate with respect to the validity of this approach (Farina and Enoka, 2011; De Luca and Nawab, 2011; Enoka, 2019; DeFreitas, 2019), and, thus, further independent examination of MU behaviour under different conditions is needed with complementary methods.

One approach to analyse and extract MU behaviour from bipolar sEMG recordings is wavelet-based analysis of frequency components. This technique was proposed by von Tscharner (2000) and has been successfully used to examine recruitment patterns of lower and higher threshold MUs in a range of applications using EMG (von Tscharner, 2002; von Tscharner & Goepfert, 2006; Wakeling & Rozitis, 2004 Hodson-Tole & Wakeling, 2007; Lee et al., 2011). Indeed, these techniques can also be applied to sEMG (Wakeling et al., 2001; Wakeling, 2009a; Wakeling, 2009b). Traditional analysis of sEMG frequency components typically considers mean/median frequency values, where all frequency components are banded together (e.g. Jenkins et al., 2015). This examination of frequency characteristics where all frequency components are banded together limits understanding of the aetiology of these mean/median frequency shifts (e.g., a reduction in mean frequency could reflect an increase in lower frequency components or a decrease in higher frequency components). In contrast, wavelet analysis quantifies signal power within defined frequency bandwidths, enabling a finer-grained assessment of the interplay between low and high frequency components, which are related to the excitation of smaller and faster motor unit populations, respectively (Hodson-Tole and Wakeling, 2009; Lee et al., 2011).

The aim of the current study, therefore, was to use a wavelet-based analysis to examine MU recruitment of the quadriceps under both higher- and lower-torque conditions (70% and 30% of MVC, respectively) using an isometric knee extension model performed to momentary failure. Specifically, we compared sEMG frequency characteristics across two continuous isometric knee extension tasks (low- and high-load) performed to momentary failure. By doing so, we sought to identify potential differences in recruitment patterns, especially at momentary failure, as determined by wavelet analysis.

## Methods

### Experimental Design

A within-session, repeated-measures, randomised crossover design was adopted to examine and compare MU recruitment patterns during isometric knee extensions with both high- and low-load demands performed to momentary failure. The study was approved by the Centre of Health, Exercise and Sport Science Research Ethics Committee (ID No. 582) meeting the ethical standards of the Helsinki declaration and was conducted within the Sport Science Laboratories at Southampton Solent University.

### Participants

A convenience sample of 10 recreationally active adult males (height: 179.6±6.0 cm; mass: 76.8±7.3 kg; age: 26±7 years) with previous resistance training experience (6±3 years) were recruited. Exclusion criteria were based upon illness or any contraindications to physical activity identified using a physical activity readiness questionnaire, though no one was excluded. All participants read a participant information sheet, were afforded the opportunity to ask any questions, and then completed informed consent forms before any testing commenced.

### Equipment

Stature was measured using a wall-mounted stadiometer (Harpenden stadiometer, Holtain Ltd, UK) and body mass was measured using balance scales (Seca 710 flat scales, UK). Trials were performed on an isokinetic dynamometer (Humac Norm, CSMi, USA). Surface electromyography was measured using a Trigno Digital Wireless sEMG System (Delsys, USA). Torque and sEMG signals were collected using the isokinetic dynamometer and sEMG systems, respectively, which were synced using a Trigno Analogue Adaptor (Delsys, USA); both torque and sEMG signals were recorded using the EMGworks Acquisition software (Delsys, USA).

### Testing

Two different conditions were examined: single continuous isometric efforts to momentary failure at 30% MVC and at 70% MVC. Both conditions were counterbalanced between participants as to whether they would perform either the 30% or 70% condition first, separated by a 20-minute rest. Prior to each condition, MVCs were performed and participants were instructed to apply maximal isometric effort against a fixed resistance at 45° of knee flexion. This procedure was completed before each condition to determine the respective absolute torque demands for each participant that equated to 30% and 70% of MVC; that is to say, each trial was normalised to the preceding MVC. Participants were instructed to gradually build up to a maximal effort over 3 seconds and were instructed to gradually reduce their effort once it was clear that a max torque had been achieved (i.e. when the torque reading was no longer increasing). In all conditions, knee angle was set at 45° flexion (0° = full extension) to standardise the exercise between participants. Before testing started, participants completed a standardised warm up of 20 body weight squats.

Each participant was instructed to perform an isometric effort with enough torque to reach their respective load for each condition. Participants were provided with a visual aid in the form of a horizontal on-screen torque bar with limits set at 70±5% and 30±5% of MVC for the high- and low-load conditions, respectively. Participants were instructed to generate enough torque to ensure a vertical on-screen torque bar was between the lower- and upper-limits until momentary failure. Participants were verbally encouraged throughout; if they fell below the lower torque limit, they were encouraged to attempt to regain their set torque output. Momentary failure was defined as when participants could no longer generate enough torque to keep within the torque limits set, despite exerting maximal effort (Steele et al., 2017).

### Surface Electromyography

Surface electromyography was recorded during each condition for the vastus medialis (VM), rectus femoris (RF) and vastus lateralis (VL). Electrode placement was made according to recommendations from the Surface Electromyography for the Non-invasive Assessment of Muscles (SENIAM) project (http://www.seniam.org/). Participants’ skin was shaved and cleaned using an alcohol-free cleansing wipe at the site used for electrode placement. Raw signals were collected at 2000 Hz.

### Wavelet Based Analysis

Myoelectric signals were decomposed into intensities as a function of time and frequency using an EMG-specific wavelet transformation (von Tscharner, 2000). A filter bank of 11 non-linearly scaled wavelets (*k*∈ {*0,1*,…,*10*}), with central frequencies (*f*_*c*_) spanning 6.90–395.44 Hz, was used. To quantify features of signal amplitude, the total intensity at each time point (*I*_*t*_) was calculated by summing the intensities over wavelets *1* ≤ *k*≤ *10*. Exclusion of the first wavelet (*k*= *0*) ensures low frequency content, associated with factors such as movement artefact, were not included in the analysis. The total intensity calculated is comparable to twice the square of the root mean square value (Wakeling et al., 2002).

To quantify changes in signal content within each frequency domain, changes in the intensity within each domain were calculated. Within each wavelet domain, an intensity (*I*_*j,k*_) was calculated for each sample point (*j*). During the trials, the intensity changed as a function of trial duration. The rate of change was calculated as the slope of the linear least squares fit of log *I*_*k*_plotted as a function of trial duration (0 to 1, representing the beginning and end of the trial, respectively). This describes the exponential change, expressed as the percentage change of *I*_*k*_per trial.

In addition, the mean signal frequency 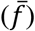 of each trial was calculated to facilitate a more direct comparison with previous literature, wherein mean or median frequency analyses are reported. This also enabled us to contrast the mean frequency results with those from our wavelet-based analyses. Here, mean frequency was calculated from wavelet transformed data (*k* = 1 – 10) using the weighted mean,

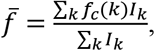

for both the start and end of the trials. The start and end of the trial was defined by torque thresholds of 25% (start) and 75% (end) of the mean torque throughout the trial. The changes in signal intensity within each frequency band were calculated between these two points so as to remove the effects of the ramping up/down of torque at the start/end of the trials. All analysis of EMG data was performed in Mathematica (version 11.1, Wolfram).

### Statistical Analysis

All code and data used for this study are available on the Open Science Framework (https://osf.io/g2z4w/). To assess the effects of load on the myoelectric signal, we performed statistical analysis in R (version 4.0.2; R Core Team, 2020). Before performing inferential analyses to answer our research question, we first visually inspected and quantified the effects of set order and condition on MVC and time to momentary failure. These were considered descriptive analyses and are presented using individual data points and mean ± SD or geometric mean ⋇ *g*eometric SD. Descriptions of trial durations and their differences were performed on the geometric (log) scale, and as such, are presented multiplicatively.

Our first analysis compared the mean frequency at momentary failure between the two loads (30 vs. 70% MVC), after adjusting for initial mean frequency. We created a linear mixed-effects model that was parameterized as an analysis of covariance,

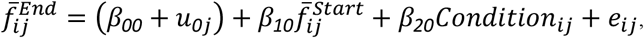

for row *i* in participant *j*, and where *u*_*0j*_ *i*s a random effect for participant *j*, 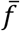 is mean frequency, and *Condition* is dummy-coded 0 for 30% and 1 for 70% MVC. Each muscle was fit in a separate model due to convergence issues when attempting to fit them together. Since the residuals were not normally distributed, compatibility intervals (CI) for the effect of condition (*β*_*20*_) were generated using the basic (reverse percentile) bootstrap with 500 replicates.

Second, we investigated the effects of load on relative changes in wavelet intensities. We created a linear mixed-effects model in which we parameterized a mean-centred log(*f*) to linearize the relative change-frequency relationship. The resulting mixed-effects model, in Pinheiro-Bates-modified Wilkinson-Rogers notation (Wilkinson and Rogers, 1973; Pinheiro and Bates, 2000) for brevity’s sake, was

`I ∼ Muscle * Freq * Condition + (Freq + Condition + Muscle | Participant)`, where `Condition` was dummy-coded 0 for 30% and 1 for 70% MVC. The mean-centring of the log(*f*) decreased collinearity between random effects, allowed for our intercepts to be interpretable, and enabled the model to converge on a solution with normally distributed and homoscedastic residuals. Thus, we parametrically calculated the CIs using estimated marginal means on the fixed effects (as a function of frequency, conditional on muscle and loading condition) and their contrasts with Satterthwaite degrees-of-freedom approximation (Lüdecke, 2018; Lenth, 2020).

Finally, we quantified the degree of wavelet similarity over time. Each high- and low-load trial was respectively summarized by a 10 (wavelets) × 1000 (time points) matrix, with which we generated a 1000×1000 concordance cross-correlation matrix for each muscle for each participant (Vigotsky, 2020). For example, a participant’s VM wavelet intensities from the 30% and 70% trials were used to generate a 1000×1000 matrix, with each element corresponding to the absolute agreement in wavelet intensities between time *i* in the 30% trial and time *j* in the 70% trial, and where agreement was quantified using Lin’s concordance correlation coefficients. We used linear mixed-effects models to summarize these concordance correlation matrices. Three models—one for each muscle—quantified the effect of trial duration on wavelet intensity agreement, wherein the percentages of trial duration of the 30% and 70% conditions were used as regressors and the concordance correlation coefficient (*ρ*^*c*^) at that time point was the response variable,

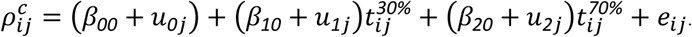

We allowed slopes and intercepts to vary and, here, we report parameter estimates (fixed-effects) along with participant-level variances (random-effects).

In addressing our research questions, we took an estimation approach rather than hypothesis-testing approach since we were neither interested in dichotomizing our findings nor comparing them to a null model (Gardener and Altman, 1986; Amrhein et al., 2019). Effects and their precision, along with the conclusions based on them, were interpreted continuously and probabilistically, considering data quality, plausibility of effect, and previous literature, all within the context of each outcome (Amrhein et al., 2019; McShane et al., 2019).

## Results

### MVC and Time to Momentary Failure

Reductions in MVC were not appreciable between the first and second trial, nor between those who completed the trials in a different order (i.e., 30% vs. 70% first) (Figure 1A). Time-to- momentary failure was, on average, 3.3 ⋇ *1*.3 -times longer (geometric mean ⋇ *g*eometric SD) for the low torque trial condition compared with the high torque trial condition. In contrast to the MVCs, there was an order effect for time-to-momentary failure. Specifically, 30% trial durations were an average of 46% longer when the 30% trial preceded the 75% trial, but 70% condition were only 26% longer when the 70% trial preceded the 30% trial (Figure 1B). Since there was an effect of order on set duration in our sample (not necessarily inferentially), we visually ensured that there was no salient effect of set duration for other outcomes. These visual checks can be found in our supplementary material.

**Figure 1.**
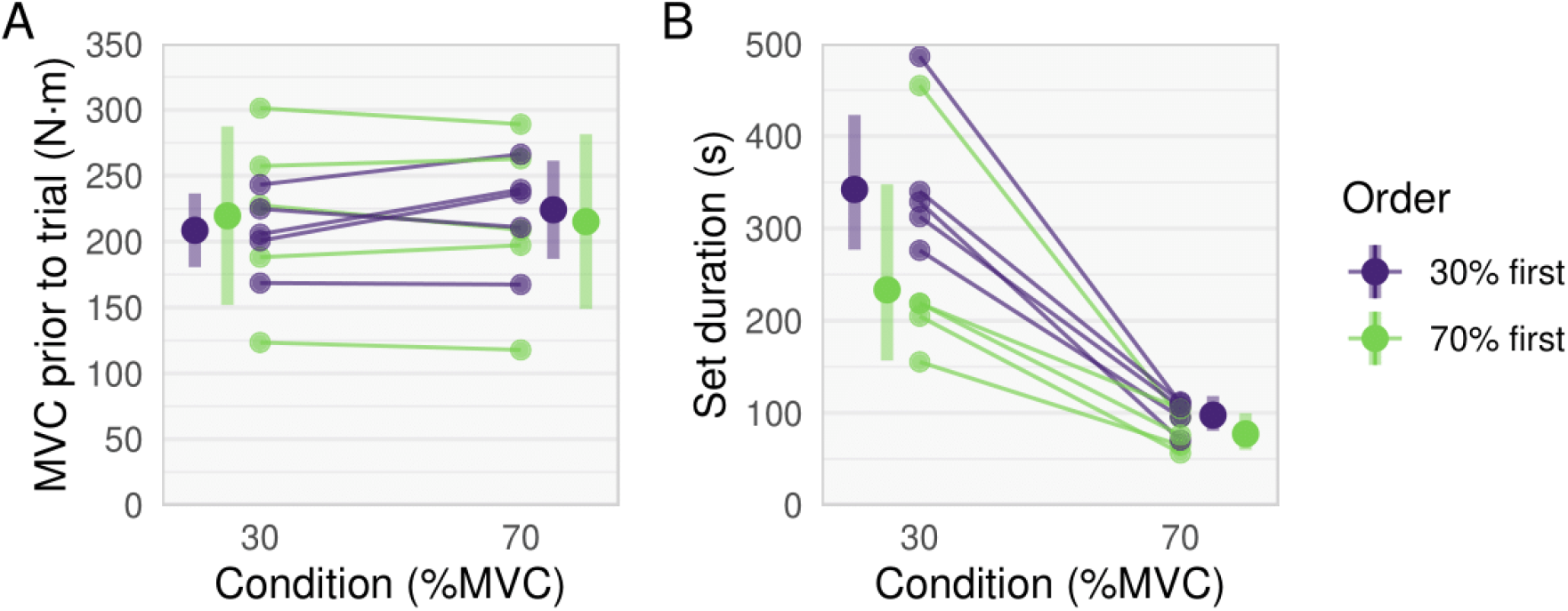
**(A)** MVC prior to trial and **(B)** set durations for each participant for the 30% and 70% conditions. Thick dots and error bars are mean ± SD in **(A)** and geometric mean ⋇ *g*eometric SD in **(B)** and participants performing the 30% condition first are shown in purple, and those performing the 70% condition first are shown in green.

### Mean Frequency

As calculated from the wavelet transformed signals, there was a decrease in mean frequency from the start to the end of both the low- and high-load trial conditions. These frequencies were similar across load conditions (Figure 2). This is supported by the ANCOVAs, which showed minimal effects of load on mean frequency at momentary failure for the rectus femoris (effect of 70% relative to 30% (CI_95%_) = 3 Hz (−14, 21)), vastus lateralis (−0.7 Hz (−21, 17)), and vastus medialis (5 Hz (−12, 25)).

**Figure 2.**
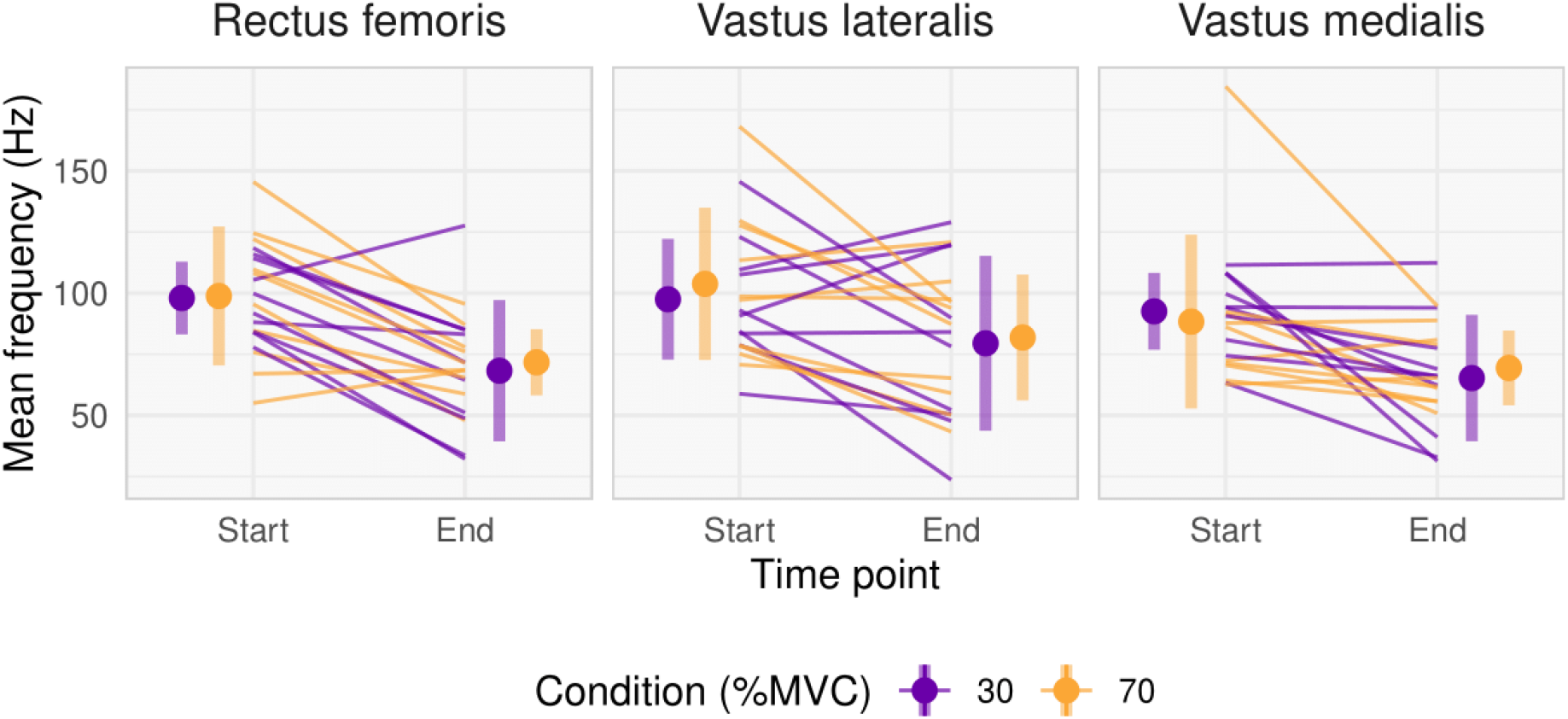
Start and End mean frequencies for the rectus femoris, vastus lateralis and vastus medialis for each participant Thick dots and error bars are mean ± SD and participants performing the 30% condition first are shown in purple, while those performing the 70% condition first are shown in orange.

### Wavelet Analysis

At the start of each muscle’s 30% condition, the greatest intensities occurred at the lower frequencies, with relatively little signal content at higher frequencies (dark blue in Figure 3). As the trial progresses, intensity increases in the low-to-mid frequency ranges while greater intensities are also shown to occur in the higher frequency components after approximately 60% of the trial duration (green-yellow in Figure 3).

**Figure 3.**
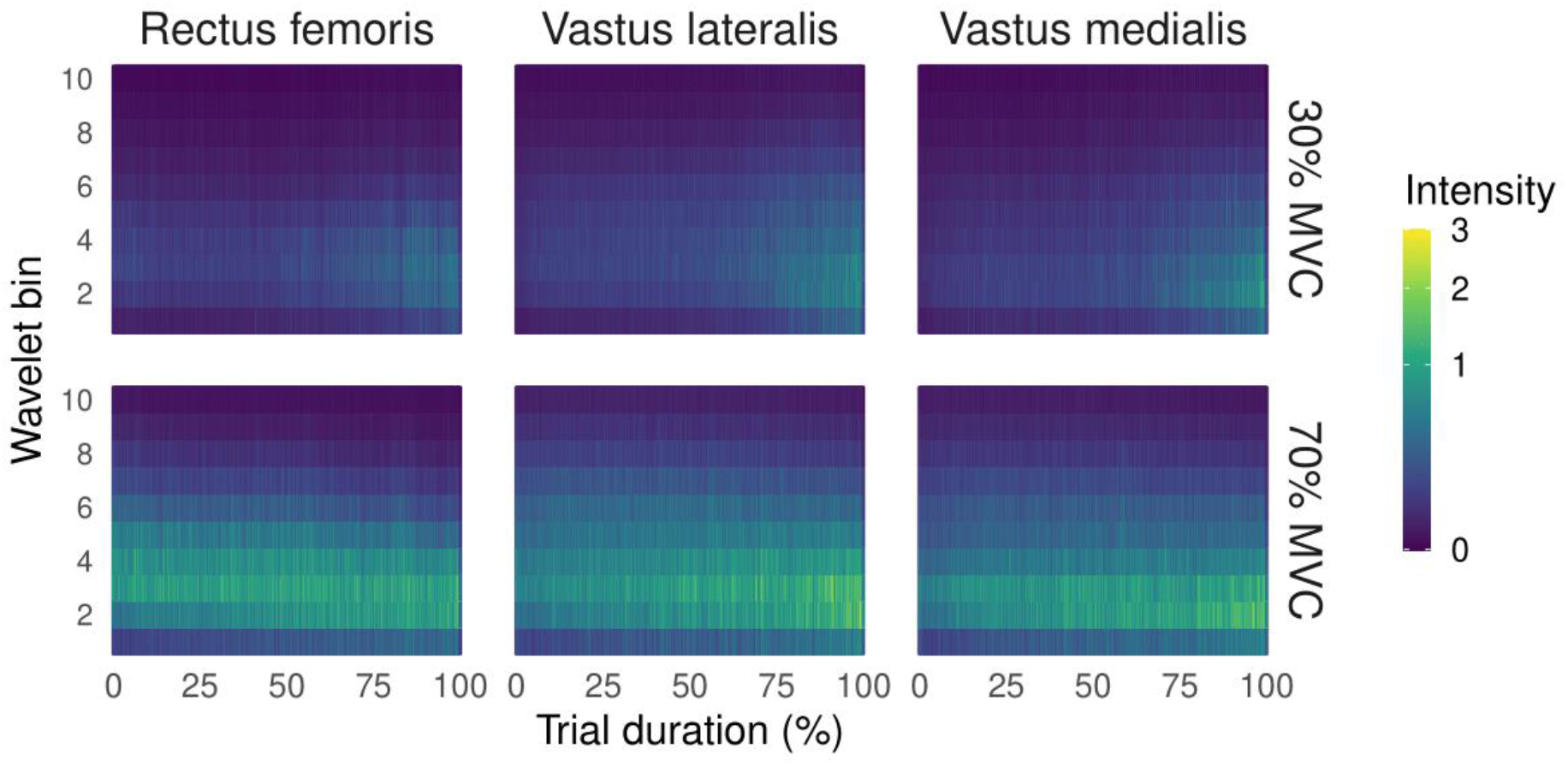
Spectrograms of wavelet intensities across time. Time is normalised to percentage of trial duration (%) across the *x*-axis; the *y*-axis indicates the wavelet bin number, with higher bin numbers corresponding to higher frequencies. Color indicates intensity of the wavelet bin at a given time point.

In the 70% condition, there is initially intensity across all frequencies (note colours extend the whole frequency spectrum in Figure 3), with the greatest intensities at mid-to-low frequencies. Over the course of the trial, there is an increase in intensity visible in the lower frequencies (note region displaying green-yellow colour in Figure 3). The reduction in intensity of the higher frequencies is not as visible in these figures but, for example, some loss of intensity can be seen in the highest wavelets at the very end of the trial, particularly in the RF.

### Wavelet Agreement

Concordance correlation coefficients increased with respect to the duration of the 30% trial and either decreased slightly (VM and VL) or remained the approximately the same (RF) with respect to the 70% trial (Table 1, Figure 4).

**Table 1.**
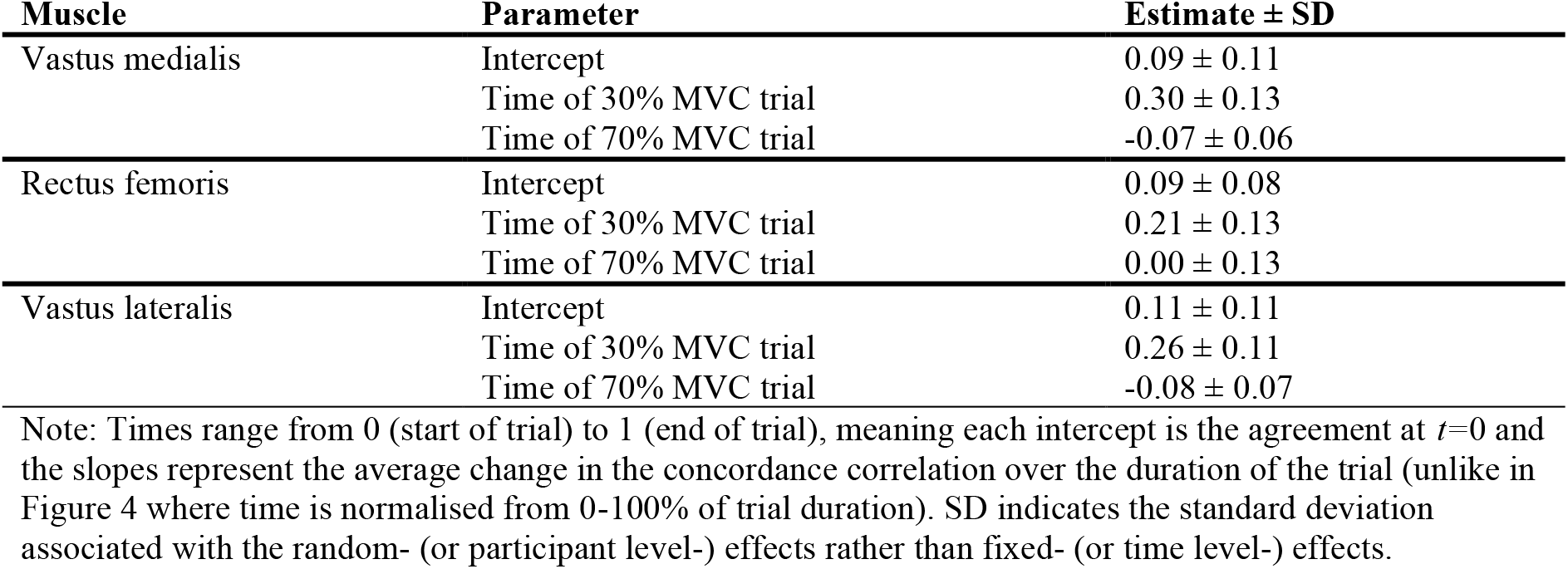
Relationship between trial durations and the agreement of wavelet intensities.

**Figure 4.**
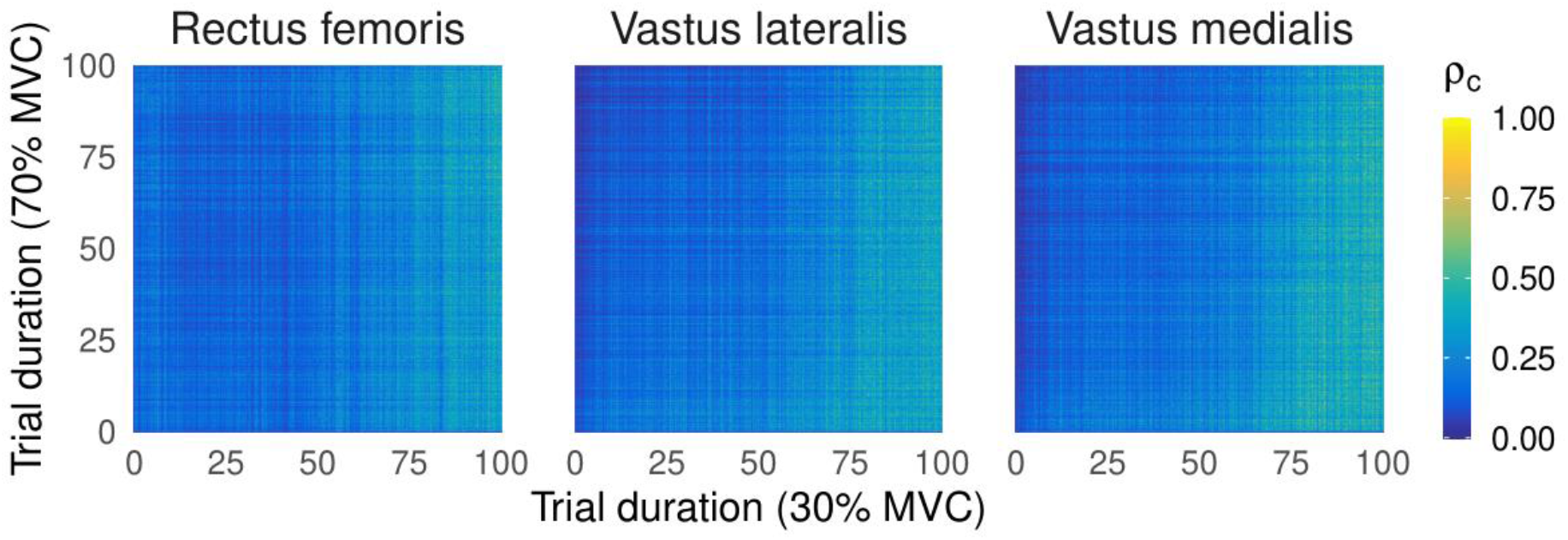
Average absolute agreement of wavelet intensity vectors across trial durations. Each point represents the average concordance correlation coefficient between the wavelet intensity vector from % of trial duration on the *x* axis for the 30% trial and % of trial duration on the y axis of the 70% trial.

### Relative Change in Signal Intensity Across Wavelets

Across all muscles, the 30% condition had greater increases in wavelet intensities, and these changes were more uniform across the frequency spectrum (Table 2, Figure 5). During the low-torque trials, it is clear there was an increase in intensity across the whole frequency spectrum studied, but these increases are larger for the lower frequencies. In contrast, during the high-torque trials, there was an increase in signal intensity of the lower frequency components, but a decrease in the intensity of the higher frequency components. The transition between increasing and decreasing occurs at different frequencies in the three muscles: ∼200 Hz in VM, ∼75 Hz in RF, and ∼150 Hz in VL.

**Table 2.**
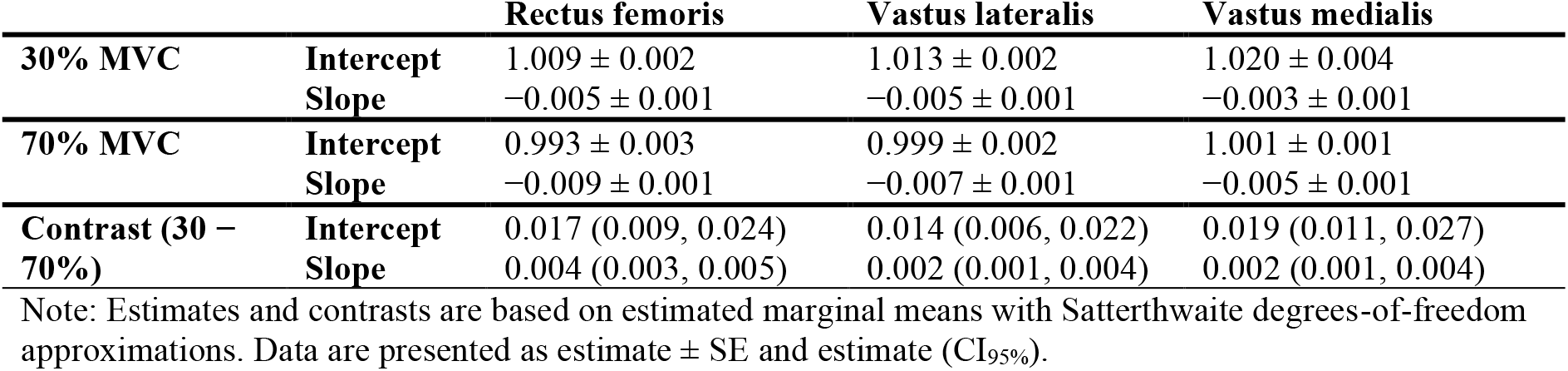
Wavelet intensity intercepts and slopes for each muscle for the 30% and 70% conditions.

**Figure 5.**
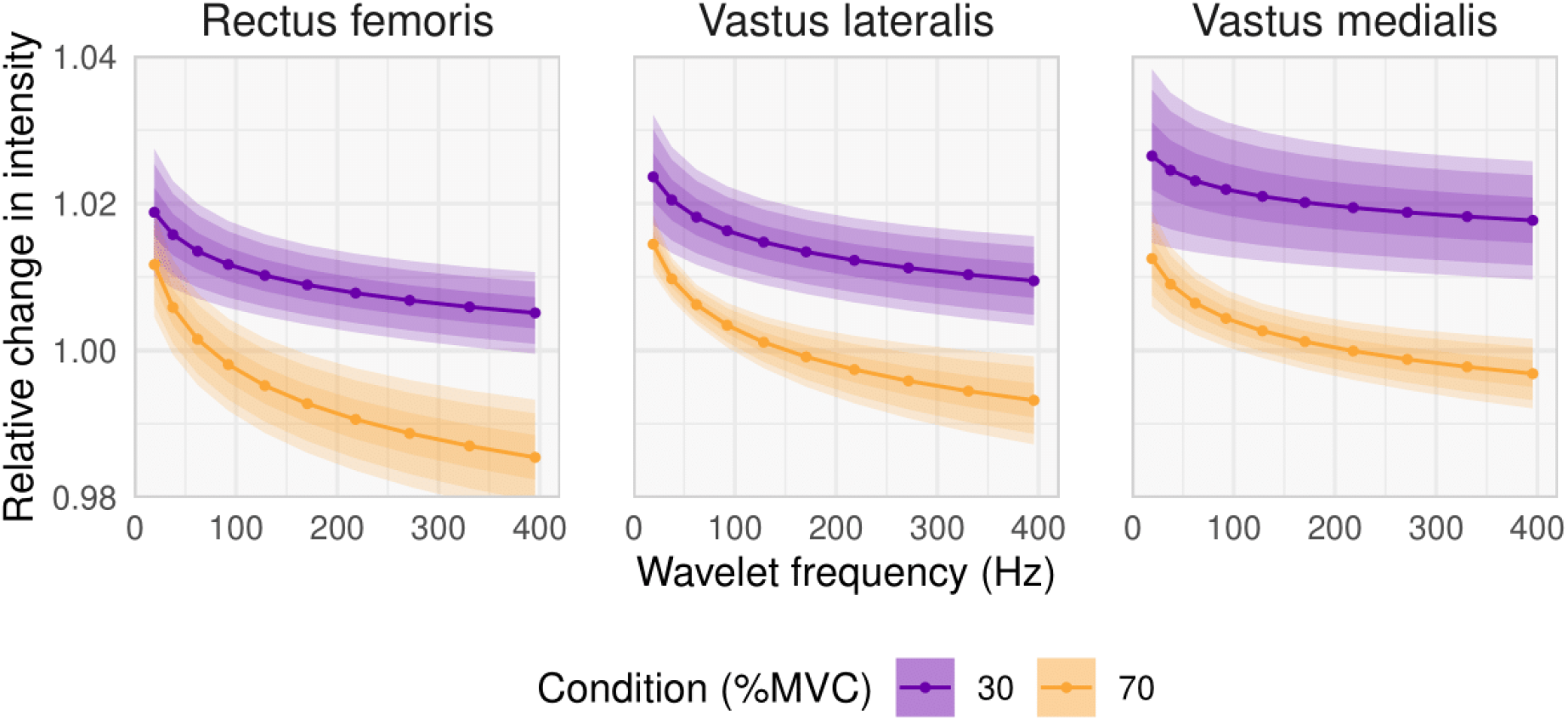
Relative changes in wavelet intensities across for each muscle for the 30% and 70% conditions. Lines and ribbons are from estimated marginal means on the model’s fixed effects with Satterthwaite degrees-of-freedom approximation. The ribbons indicate CIs at the 68, 95, and 99% level, corresponding to approximately 1, 2, and 3 standard errors.

## Discussion

This appears to be the first study to examine MU recruitment patterns *in vivo* using wavelet-based analysis of sEMG under both high- and low-torque conditions to momentary failure. Wavelet-based calculation of the mean signal frequency appeared to show similar mean frequency characteristics when reaching momentary failure. However, inspection of the individual wavelets reveals that changes differed between frequency components, suggesting that patterns of recruitment may have differed. Low-torque conditions resulted in an increase in intensity of all frequency components across the trials for each muscle (Figures. 2-3). This suggests evidence of additional MU recruitment (both lower and higher threshold MUs), likely to maintain torque production throughout the task. In contrast, high-torque conditions resulted in a wider range of frequency components contained within the myoelectric signals at the beginning of the trials (Figure. 3), suggesting an initial recruitment of a wider range of MUs. However, as the low-torque trial neared momentary failure there was an increased agreement between conditions across wavelets (Figure. 4). In addition, there was an increase in the signal intensity within low- to mid-frequencies over the trial while a reduction in higher frequency components occurred (Figure. 5). This potentially suggests firing rate adaptations or changes in conduction velocity occurred in higher threshold MUs as the trial progressed. These findings are also in line with those expected from recent modelling (Potvin and Fuglevand, 2017)^1^.

The differences in signal intensity changes within the different frequency components, between the high- and low-torque conditions, is quite striking (Figure. 5). In both cases, there is a relatively greater increase in the low frequency content compared to high frequencies. This increase could be representative of several factors, including MU recruitment. However, this must be interpreted with caution as these changes could equally result from fatigue-related changes in membrane properties that would influence both action potential shape and conduction velocity and, therefore, the signal frequency components (Mortimer et al., 1970; Brody et al., 1991; Dimitrova and Dimitrov 2003; Dideriksen et al., 2011). Given that the increase in lower frequency components is common to both high- and low-torque trials, we can only speculate as to the cause of these changes.

In contrast, relative changes in wavelet intensities differed between conditions. These differences were especially apparent in the higher frequency components in each of the three muscles. An increase in relative change across all frequencies was a common feature of each of the three muscles for the low torque trial, while intensity reductions within the higher frequencies occurred for high torque trials (Figure. 3). The fact that opposite effects were observed between the two conditions indicates that different myoelectric signal properties resulted from the two conditions. The increases seen during the low-torque trials could be indicative of recruitment of larger high-threshold MU populations, which may have occurred in response to the growing fatigue. The reduction in the intensity, within the higher frequency components during the high-torque trials, could be indicative of reduced recruitment of these populations of larger motor units, potentially resulting from them becoming fatigued. Alternatively, this latter effect in the high torque conditions could also be due to changes in action potential shape and conduction velocity from fatigue (Mortimer et al., 1970; Brody et al., 1991) or firing rate adaptations as previously mentioned (Potvin & Fuglevand, 2017).

Our analyses of different frequency components revealed different recruitment patterns across the conditions, which could not be identified through amplitude or mean frequency analyses alone. As a point of comparison with our wavelet analyses and with other literature (e.g. Jenkins et al., 2015), as mean frequency is commonly used as a crude assessment of MU recruitment (Phinyomark et al, 2012), we examined the mean frequency across the individual wavelets (Ranniger and Akin, 1997). In the present study, we found that mean frequency decreased similarly across both conditions and muscles. However, our wavelet-based analyses provide further insight into why such mean frequency changes occur which, in this case, were likely a result of the increases in lower frequency components within the signal (Figure. 3). Thus, wavelet-based analysis potentially offers greater insight into how the patterns of recruitment differ between conditions as a trial progresses towards momentary failure, and indeed upon reaching momentary failure there was largely similar frequency content between conditions in our study (Figures. 3 & 4)

Our findings are in contrast to the results of recent work using decomposition sEMG. Muddle et al. (2018) reported that, upon reaching momentary failure, the higher-torque condition typically resulted in greater recruitment of higher threshold MUs on average. The seemingly discrepant results may be due to the nature of the analyses performed. With decomposition sEMG, it is possible to examine the individual firing trains of MUs (4670 MUs in total in the case of Muddle et al.; i.e., ∼88 MUs per participant for high torque and ∼171 for low torque), whereas the use of wavelet analysis of bipolar electrodes permits the examination of MU ‘pools’ based upon their frequency characteristics. Thus, the former allows examination of individual MU recruitment patterns, and the latter permits examination more broadly of the recruitment patterns of populations of MUs within the muscle.

Further, although the same relative torques were used (30% and 70% of MVC), Muddle et al. (2018) had participants perform repeated trapezoidal isometric muscle actions whereas we had participants perform a single continuous isometric action. Motor unit recruitment strategies likely differ between dynamic and isometric conditions (van Bolhuis et al., 1997; Babault et al., 2006) so they may also differ between repeated isometric muscle actions and continuous efforts. For example, metabolite increases are greater during continuous isometric actions compared with intermittent (Schott et al., 1995) so this might influence recruitment, or at least the signal properties and, thus, the results of analyses and their interpretation. The results of the present study are similar with those of Muddle et al. (2018) in that there appeared to be increased MU recruitment across both high- and low-torque conditions. Indeed, Harmon et al. (2021) also recently reported that recruitment of MUs increased during low-torque conditions and, at momentary failure, were like that observed during non-failure high-torque conditions. Our study revealed increases in the signal frequency components of the individual wavelets, suggesting increased recruitment in both conditions, yet with seemingly different patterns of recruitment between conditions.

With the high-torque condition, though there were increases in lower frequency signal components, there were also reductions in the higher frequency components. This is also perhaps to be expected as Potvin and Fuglevand (2017) found in their modelling study that loss of force capacity was always higher in the upper to middle range of MUs. Upon starting the conditions, it is likely that the higher-torque condition necessitated the recruitment of more, and indeed larger, MUs compared with the lower-torque condition (Henneman, 1957). As such, the reduction in these higher frequency components in the higher-torque condition may reflect their fatigue, reduced excitation, changes in conduction velocity, de-recruitment, and/or firing rate adaptations.

It could be argued that our results suggest additional MUs are being recruited in the face of insidious fatigue in both conditions and that the pools of MUs recruited across the task, and upon momentary failure, may be similar. Yet, subtle differences in recruitment patterns appear to occur dependent upon the torque requirements of the conditions. These findings dispute the conclusions drawn by other investigators interpreting higher sEMG amplitudes as indicating greater MU recruitment with higher-loads. Further, these results potentially lend support to the explanations offered regarding MU recruitment to explain the similar adaptations that occur between resistance exercises with high- and low-loads performed to momentary failure (Carpinelli, 2008; Fisher et al., 2017; Potvin and Fuglevand, 2017; Morton et al., 2019).

Despite our efforts to strictly control our experiment, and perform a rigorous analysis, our findings may still have been affected by uncontrolled natural factors that affect the sEMG signal. Specifically, the signal frequency content of sEMG is affected by many factors (muscle fibre depth and length, pennation angle, cellular environment, temperature, fatigue, motor unit synchronisation). We were able to control for several of these factors by collecting data on the same day, from the same individuals, with the same electrode placement and joint angle configuration during an isometric effort. Thus, factors relating to muscle anatomy (i.e., muscle fibre depth and length) would have been unlikely to contribute to any differences; though it is possible that small changes in pennation angle may have occurred due to tendon creep within trials. Further, although muscle temperature was not measured, participants completed all trials using the same procedures, and in a randomised counterbalanced fashion, so this did not likely contribute to differences between conditions.

As mentioned, fatigue can lead to a reduction in signal frequency content, yet our data showed distinct patterns between loads which were similar across muscles, thus indicating there are different physiological processes underlying the signal, including differences in MU recruitment strategies. MU synchronisation, such as when generating forces rapidly, can also lead to alterations in the spectral profile of the myoelectric signals. This phenomenon typically leads to a spike in the 30 Hz region. Thus, the ‘synchronous’ MU recruitment that appears to have occurred initially in the high-load condition may have been influenced by this effect. However, the initial rate of torque production at the beginning of the trial did not differ substantially between the two load conditions so firing rate could be expected to be similar in each. Further, we used defined torque threshold to exclude from analysis the periods of each trial where ramping up/down of torque was occurring at the start/end. Additionally, when analyses were re-run with wavelets, but with the 30Hz data discarded (rather than just the first wavelet), it made no difference to our results. As such, it seems the differences in the results do not reflect differences related to firing rate modulations borne out during early ramping, or late ramping down, of torque production during trials.

## Conclusions

Wavelet-based analysis of MU recruitment patterns suggest that an increase in recruitment occurs during both high- and low-torque isometric conditions. Importantly, similar MUs appear to be recruited in both conditions upon reaching momentary failure as indicated by the similarity in frequency content at momentary failure. However, patterns of recruitment differ between conditions with high-torque conditions showing greater initial ‘synchronous’ MU recruitment, yet reduction in high frequency signal components as trials progressed towards momentary failure and, thus, fatigue of some larger MUs. Low-load conditions demonstrated increases across all frequency components of the signal, suggesting ‘sequential’ MU recruitment. Thus, when performing isometric efforts to momentary failure, our data corroborate modelling studies as well as recent biopsy evidence, suggesting overall MU recruitment may largely be similar with the highest threshold MUs likely recruited, despite being achieved with differences in the pattern of recruitment over time utilised.

## Footnotes

1 Indeed, for comparison we have rerun the model from Potvin and Fuglevand (2017) using the 30% and 70% conditions used in the present study and include these in the supplementary materials. We show the adapted MU strength and firing rates over time for the whole trials and the final 30% of time for the 30% condition trial (supplementary file https://osf.io/9jznr/).

